# cellsnake: a user-friendly tool for single-cell RNA sequencing analysis

**DOI:** 10.1101/2023.05.03.539204

**Authors:** Sinan U. Umu, Karoline Rapp Vander-Elst, Victoria T. Karlsen, Manto Chouliara, Espen Sønderaal Bækkevold, Frode Lars Jahnsen, Diana Domanska

## Abstract

**Background:** Single-cell RNA sequencing (scRNA-seq) provides high-resolution transcriptome data to understand the heterogeneity of cell populations at the single-cell level. The analysis of scRNA-seq data requires the utilization of numerous computational tools. However, non-expert users usually experience installation issues, a lack of critical functionality or batch analysis modes, and the steep learning curves of existing pipelines.

**Results:** We have developed cellsnake, a comprehensive, reproducible, and accessible single-cell data analysis workflow, to overcome these problems. Cellsnake offers advanced features for standard users and facilitates downstream analyses in both R and Python environments. It is also designed for easy integration into existing workflows, allowing for rapid analyses of multiple samples.

**Conclusion:** As an open-source tool, cellsnake is accessible through Bioconda, PyPi, Docker, and GitHub, making it a cost-effective and user-friendly option for researchers. By using cellsnake, researchers can streamline the analysis of scRNA-seq data and gain insights into the complex biology of single cells.

## Background

Single-cell RNA sequencing (scRNA-seq) is a method used to study gene expression at the single-cell level. This stands in contrast to bulk RNA sequencing, which provides information only on the average transcript expression within a population of cells. With recent technological advancements and decreasing sequencing costs, scRNA-seq has become increasingly accessible, enabling researchers to identify novel cell types, cell states, and cellular interactions [1–4].

A standard scRNA-seq bioinformatics workflow typically involves several steps, including data filtering, normalization, scaling, dimensionality reduction, clustering, visualization, differential expression analysis, functional analysis, and annotation [4,5]. Various analysis workflows for different platforms (i.e. 10x Genomics, Drop-seq, inDrops, SMART-seq2, and Fluidigm C1) have been developed to process, analyze and holistically visualize scRNA-seq data [2,6–8]. Popular workflows like Seurat [9], SingleCellExperiment (of Bioconductor) [7], and Scanpy [6] have extensive features for scRNA analysis. The analysis of scRNA-seq data poses several challenges, including the high-dimensional data structure, technical issues (e.g. dead cells, doublets, and low unique molecular identifier (UMI) counts), batch effects, low expression levels, and the presence of complex cell subsets with multiple cell states [5]. To address these, a variety of supplementary bioinformatics tools have been developed. While some of these can be integrated into existing workflows, many require substantial expertise and bioinformatics knowledge.

Another challenge is working with multiple scRNA-seq datasets. Comprehensive documentation for the analysis of a single sample using recommended parameters is usually provided. However, it is hard for a regular user to keep track of all the decisions taken during analyses, especially if more than one sample is available. This also creates challenges if one wants to see the effect of basic parameter changes and document the results for further hypothesis testing. It is also challenging to harness the power of high-performance computing (HPC) systems when needed. There are some efforts to make batch analysis, such as the cloud-based system SingleCAnalyzer [10], the R package scTyper [11], the web application Cellenics (open-source software of Biomage), and Single-Cell Omics workbench on Galaxy (singlecell.usegalaxy.eu). Cellranger from 10x genomics also provides dataset clustering and basic differential expression analysis [12] for initial quality control (QC). However, all these workflows have limited functionality or were designed for a specific need. Online (or cluster-based) solutions might also not be suitable due to data privacy rules for sensitive data or do not provide compatible files (e.g. R data files) for downstream analysis on another platform.

Here, we introduce cellsnake, a platform-independent command-line application and pipeline for scRNA-seq analysis. Cellsnake provides a reproducible, flexible, and accessible solution for most scRNA-seq data analysis applications. One of the key features of cellsnake is its ability to utilize different scRNA-seq algorithms to simplify tasks such as automatic mitochondrial (MT) gene trimming, selection of optimal clustering resolution, doublet filtering, visualization of marker genes, enrichment analysis, and pathway analysis. Cellsnake also allows parallelization and readily utilizes high-performance computing (HPC) platforms. In addition to that, cellsnake provides metagenome analysis if unmapped reads are available. Another advantage of cellsnake is its ability to generate intermediate files (such as R data files) that can be stored, extracted, shared, or used later for more advanced analyses or for reproducibility purposes. With cellsnake, researchers can perform scRNA-seq data analysis in a reproducible and efficient manner, without requiring extensive bioinformatics expertise.

## Methods

### Cellsnake workflow and tools

The cellsnake wrapper was written in Python, while the main workflow was implemented in Snakemake [13]. To find optimal cluster resolution, we utilized clustree [14]. Seurat analysis pipeline [8] provides all the main functions required for processing scRNA data in cellsnake. These functions are wrapped into different R scripts which can also be used as standalone scripts by advanced users. Cellsnake facilitates automatic format conversion when required. For instance, CellTypist [15] requires AnnData format, and the workflow converts the files back to the required file format in R. By default, cellsnake stores files into two folders: analyses and results. The analyses folder contains metadata and R data files, which can be accessed by the user. Seurat is used for integration, and after integration, the workflow runs on the integrated dataset automatically, and the output files are stored in separate folders (i.e. analyses_integrated and results_integrated).

### Parameter selection and autodetection

Cellsnake provides Seurat’s default values for fundamental parameters like min.cells (i.e features detected at least this many cells) or min.features (i.e. cells at least this many features). In addition, non-default parameters can be provided using a YAML file, and a YAML file template can be printed and edited. Cellsnake determines which principal component exhibits cumulative percent greater than 90% and % variation associated with the principal component as less than 5 (as described hbctraining.github.io/scRNA-seq/lessons/elbow_plot_metric.html). To filter MT genes, cellsnake uses the miQC tool [16]. If that fails, it uses the median absolute deviation of the MT gene expression as an alternative. MultiK algorithm [17] is used to determine optimal resolution detection and doublet filtering is done using the DoubletFinder tool [18]. Autodetection of parameters is not offered as a default option in cellsnake due to its computational expense and potential for failure with large sample sizes. Cellsnake utilizes a special directory structure for MT percentage and resolution the results will be saved in different folders named after the selected parameters. These results are not overwritten and can be reviewed later, or the parameters can be modified for further investigation.

### Cellsnake testing and benchmarks

To test cellsnake, we obtained four samples containing exclusively macrophages from gut mucosal tissue [19], along with two fetal brain datasets [20] and six fetal liver datasets [21]. The fetal brain datasets were provided in matrix file format, while the other datasets were in FASTQ format and were processed by Cellranger (v.7.0.0) with the default settings and the default databases. For a comprehensive evaluation, we compared the features of Cellsnake with two other holistic tools, Cellenics and Single Cell Omics workbench (https://singlecell.usegalaxy.eu/). The Cellenics community instance (https://scp.biomage.net/) is hosted by Biomage (https://biomage.net/).

## Results

### Cellsnake can be run either as a Snakemake workflow or as a standalone tool

Cellsnake utilizes a variety of tools and algorithms (Table 1) and consists of two primary components: the main workflow and the wrapper. The cellsnake wrapper assists with the main workflow and provides an easy-to-use option for users. The workflow (Fig. 1) is primarily designed using the Seurat pipeline (v4.2) and the Snakemake workflow manager. As needed, the workflow integrates various algorithms to enhance the basic functionality of Seurat. For instance, when one droplet encapsulates more than one cell, it appears as a single cell and can affect the downstream analysis. Addressing this issue in the workflow is crucial [22]. A distinctive feature of cellsnake is its default doublet filtering option, a functionality not included in the standard Seurat pipeline. Users can also adjust other parameters by modifying the configuration files, which are formatted in YAML. This flexibility empowers precise analysis of scRNA-seq data.

**Figure 1.**
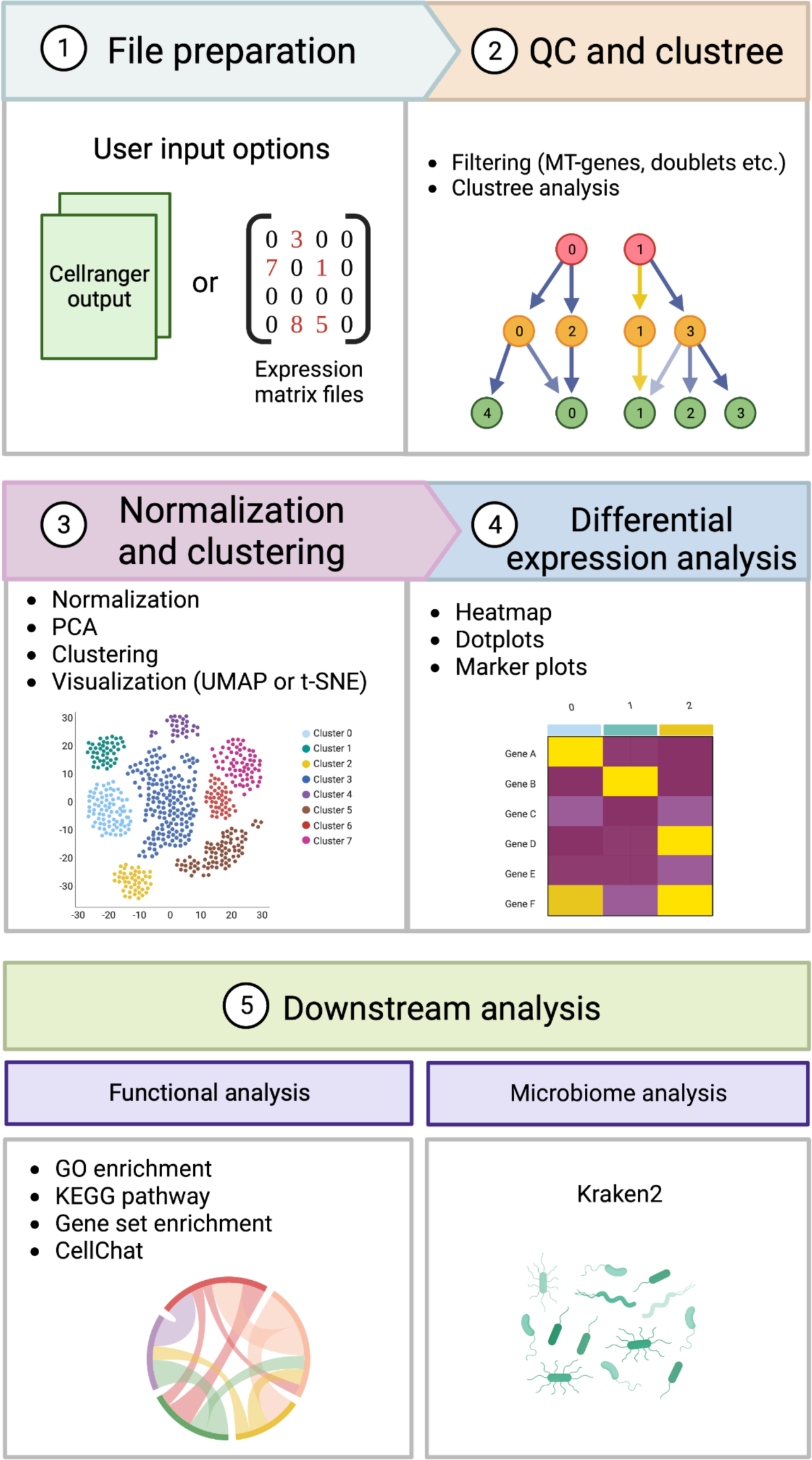
Overview of the scRNA-seq pipeline in cellsnake. (1) Cellsnake can accept the output files from Cellranger in addition to raw expression matrix files if provided in an appropriate format. (2) QC is performed by filtering out MT-genes, doublets, and cells with a low gene number as examples. Clustree is then used to find the optimal resolution for the dimensionality reduction. (3) Afterward, the dataset is normalized and scaled before the PCA analysis and visualized by UMAP or tSNE. (4) To find the differences in gene expression levels within the dataset differential gene expression analysis is performed with several outputs such as heatmaps, dot plots, and marker plots. (5) To get an even better insight into the dataset, the pipeline contains several functional analyses such as GO enrichment, KEGG pathway, gene set enrichment, and CellChat. Metagenome analysis is also available if the input file from step 1 is the direct output from Cellranger. This is done by using the metagenomics tool Kraken2.

**Table 1.**
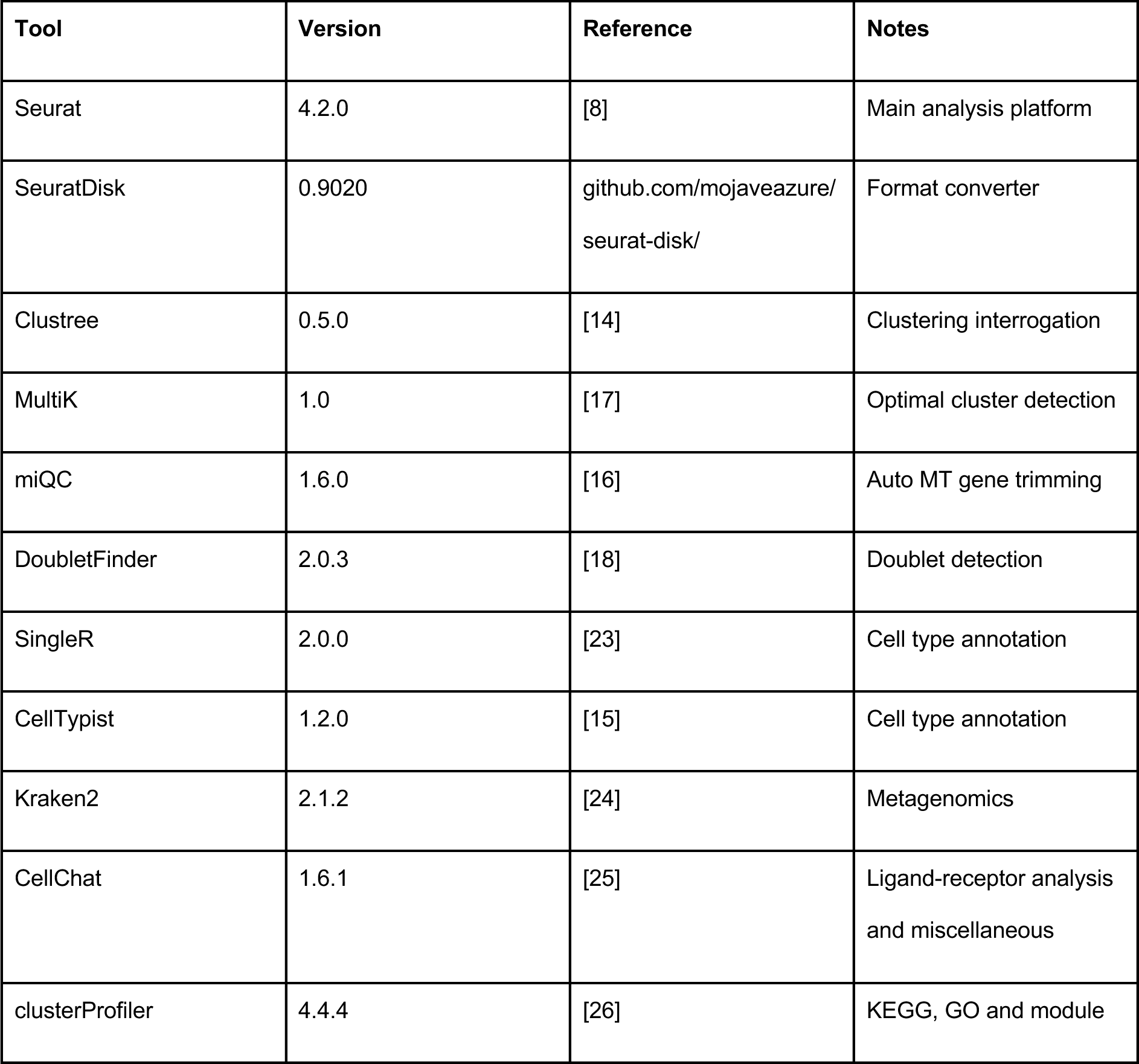
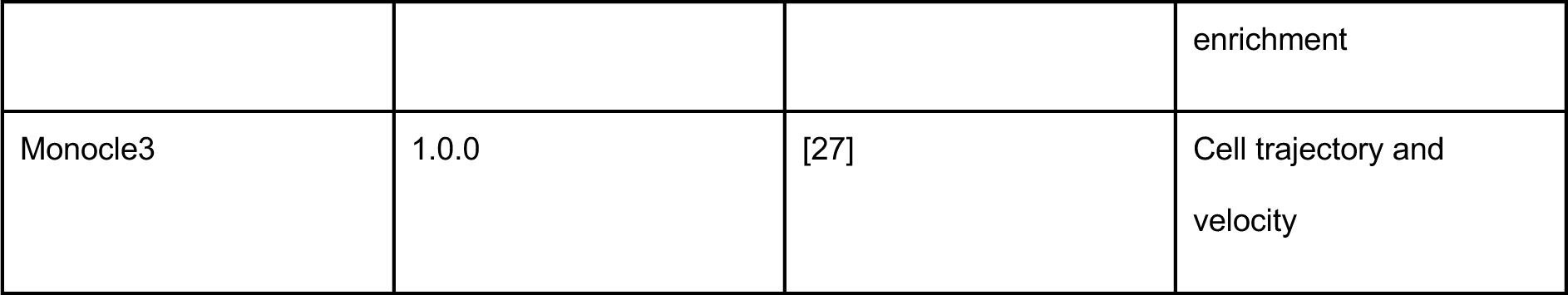
An overview of the tools and algorithms used in the cellsnake workflow, as well as an explanation of what they do and which versions are used.

Cellsnake covers most of the methods offered by Seurat, including integration. The workflow is automatically repeated for an integrated dataset once the analyses have been concluded for all individual samples available in the study (i.e QC, filtering etc.). Analysis outcomes such as dimension reduction, clustering, differential expression analysis, functional enrichment, and cell type annotations are reported for the integrated sample. Since the datasets individually passed the initial QC and are trimmed for artifacts, these steps are skipped by the workflow. Cellsnake can also generate publication-ready plots for both individual and integrated samples. It also automatically produces plots for markers (i.e. genes) which can be investigated to better understand the predicted clusters (i.e. cell subsets). Additionally, cellsnake provides the option to produce supplementary plots, featuring dimension reduction and expression images for selected genes or markers. This functionality adds a valuable level of customization to the analysis, enabling the user to explore targeted genes or markers of interest in greater detail.

The input of cellsnake can be either Cellranger output directories for batch analysis or single expression matrix files (e.g. h5 files) for individual sample processing. Cellsnake automatically detects the input format and runs accordingly with minimal user intervention and with minimal lines of input commands (Table 2). The Cellsnake workflow offers three primary modes with distinct options: minimal, standard, and advanced. The minimal mode is suitable for fast analysis, parameter selection, and downstream integration. Fundamental parameters, such as filtering thresholds and clustering resolution can be determined via a minimal run at an early stage which will reduce computational cost. Standard and advanced workflow modes contain additional features and algorithms (Table 2).

**Table 2.** Cellsnake commands and a summary of their outputs.

| Mode | Outputs | How to run? |
| --- | --- | --- |
| cellsnake minimal | Dimension reduction plots, QC metrics, technical plots (MT, counts, gene, feature), clustree plot | `$ cellsnake minimal data`<br>OR<br>`$ snakemake -j 5 --config option=minimal` |
| cellsnake standard | All minimal outputs and CellTypist, singleR annotations, enrichment analyses tables, trajectory plots, summarized marker plots | `$ cellsnake standard data`<br>OR<br>`$ snakemake -j 5 --config option=standard` |
| cellsnake advanced | All standard outputs and CellChat results, detailed top markers per cluster plots | `$ cellsnake advanced data`<br>OR<br>`$ snakemake -j 5 --config option=advanced` |
| cellsnake integrate | A single integrated object for analysis. | `$ cellsnake integrate data`<br>OR<br>`$ snakemake -j 5 --config option=integrate` |

\* data folder may contain multiple samples and this will trigger a batch analysis.

### Reanalyses of publicly available datasets using cellsnake

We showcase some features of the pipeline using publicly available datasets. The first dataset is from the fetal brain containing (only) count tables from two samples (Fig. 2 and Fig. 3). We processed two samples using the default settings (e.g. MT filtering threshold 10 percent and resolution parameter 0.8). Minimal mode only takes four minutes in a laptop for two samples of the fetal brain dataset. Another five minutes is enough for both integration and processing of the integrated sample with minimal mode. The user can decide on the parameters early on (Fig. 2) and the standard mode will finish in 50 minutes without parallel processing. Cellsnake utilizes different tools and provides outputs for all as figures (supplementary figures 1-6) or as tables.

**Figure 2.**
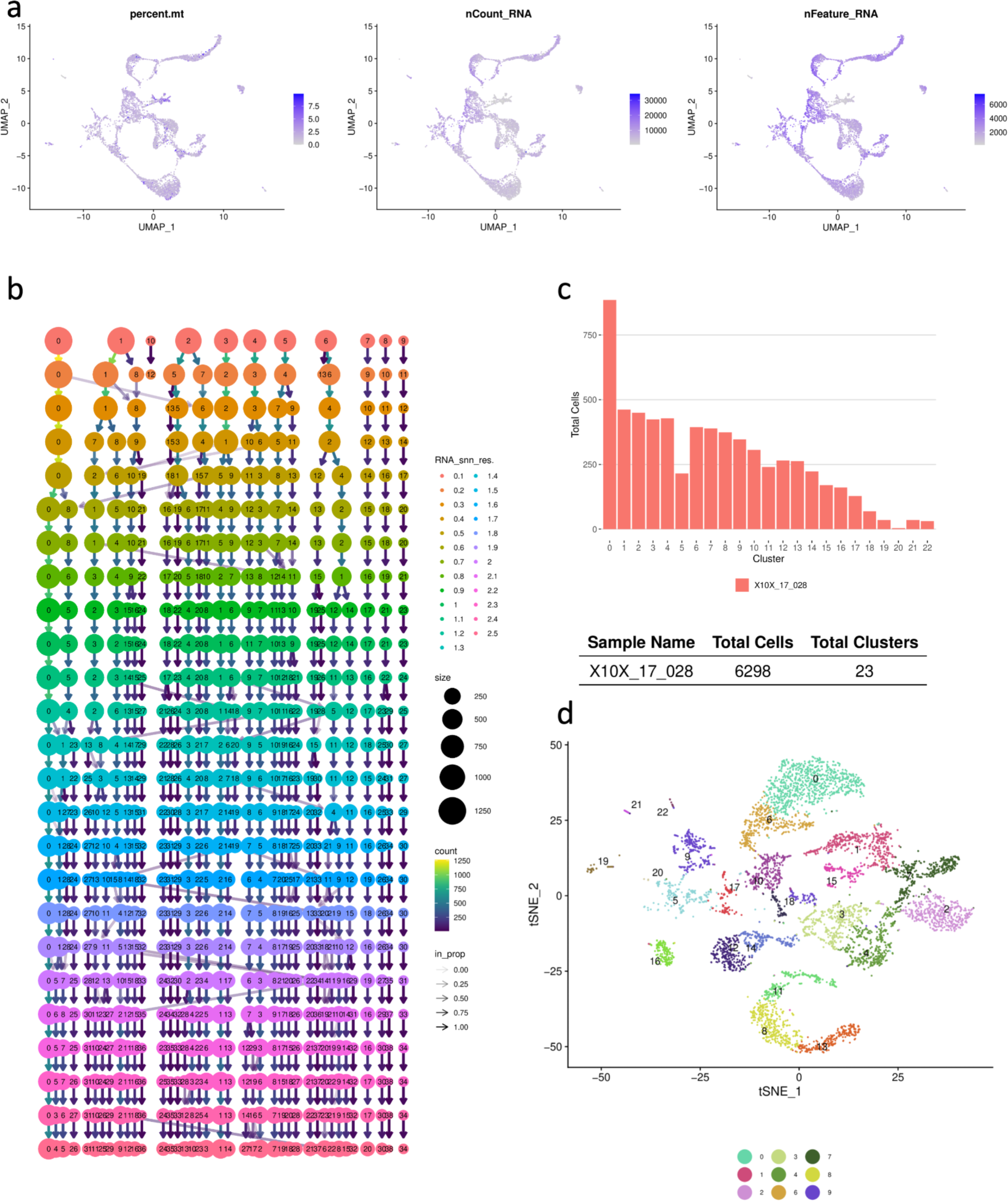
Cellsnake quickly generates standard output plots that include technical information. a) The user can investigate the fundamental statistics like MT gene percentage, number of genes detected, and reads mapped per cell information. Here the results shown are based on one of the fetal brain samples. b) Clustree analysis is not part of the Seurat pipeline but cellsnake offers this by default. This plot can be used to find the optimal number of clusters. c) The selected resolution resulted in 23 clusters and 6298 cells passed the filtering thresholds (after filtering doublets and low-quality cells). d) tSNE plot shows the clusters. Cellsnake prints only the top clusters in the legend to prevent overplotting. The user will get UMAP, PCA, and tSNE plots by default.

**Figure 3.**
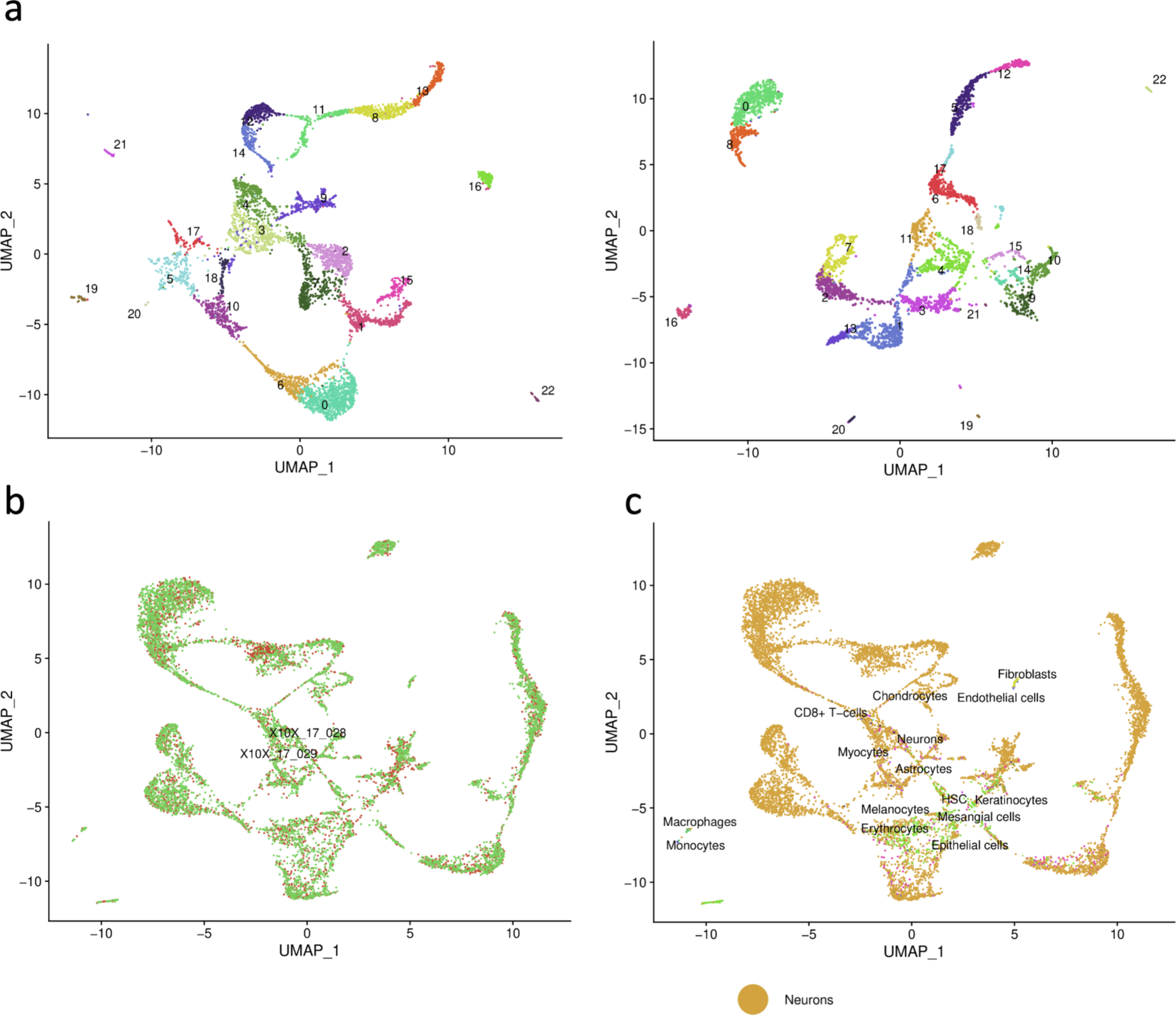
Cellsnake processes integrated samples similar to the individual samples and generates the same plots. a) The UMAP plots were generated for two samples from the fetal brain dataset, seen in the first and second panels. b) The UMAP plot shows clusters for the integrated samples. c) The UMAP plot shows cluster annotation based on the singleR package “BlueprintEncodeData” model predictions. The results showed the cells were mostly predicted as neurons which are consistent with the dataset but there are also some mispredictions. The detailed annotations can be accessed as Excel tables and heatmaps.

The second dataset is from the fetal liver containing three CD45+ and three CD45-FACS sorted samples from three different donors (Fig. 4A). This time we selected automatic filtering of MT gene abundant cells rather than a hard cut-off when pre-processing the samples. In total, 29045 cells passed the filtering threshold. The standard workflow took 3 hours with only two CPU cores on a standard laptop, which is enough for most use cases. The samples were later integrated and the optimal number of clusters was predicted automatically. The separation of two groups (Fig. 4A and 4B) in the integrated dataset is similar to what was reported in the original study [21], which indicates that cellsnake is capable of reproducing key findings from published studies. The differential expression analysis also reveals that the AHSP gene is highly expressed in CD45+ samples, which is in line with the known function of this gene in erythroid cells.

**Figure 4.**
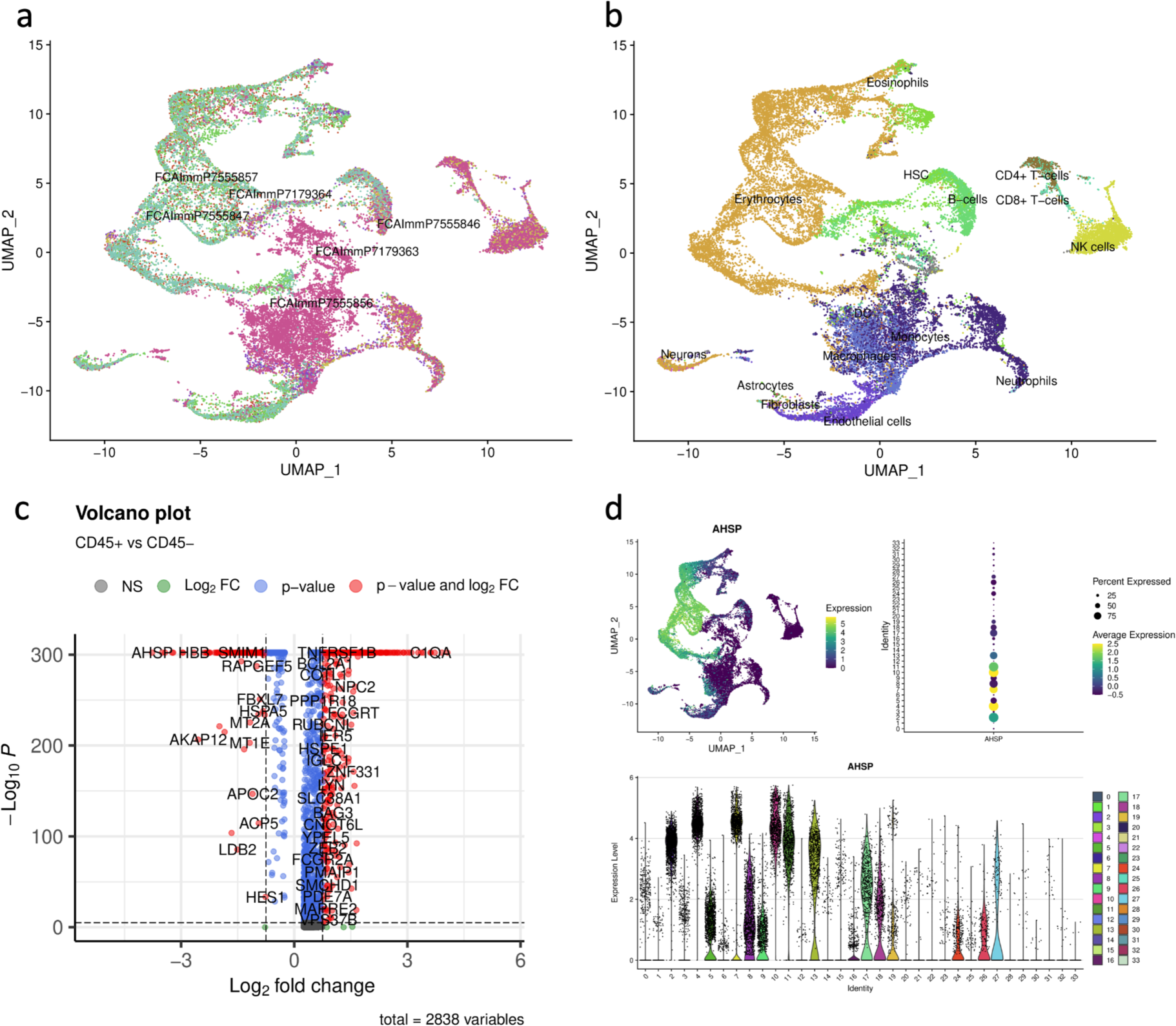
The fetal liver dataset consists of six FACS-sorted samples, integrated by cellsnake. a) Cellsnake displays integrated UMAP plot, and labels, and b) annotates the clusters. c) The user can provide the clinical information which shows differentially expressed genes among two groups. d) It is also possible to visualize selected marker genes. For example, the AHSP gene is upregulated in CD45+ samples compared to CD45-samples.

### Cellsnake can analyze metagenomics from single-cell data

Another unique feature of Cellsnake is its ability to perform metagenomics analysis using Kraken2. If a database is provided, Cellsnake will automatically run Kraken2. After collapsing read counts to a taxonomic level based on user input, such as genus or phylum, results are reported accordingly. Cellsnake provides metagenomic results in the form of dimension reduction plots and barplots, and users can load metadata into R for personalized downstream analysis.

This feature was tested on four samples from mucosal macrophages, with automatic trimming of MT genes and selection of resolution (Fig. 5). Cellsnake reported results based on the optimal number of clusters, and non-human material detected by Kraken2 is visualized on integrated UMAP plots (Fig. 5a,b). Users can also obtain a detailed list of results based on the selected taxonomic level in an Excel file.

**Figure 5.**
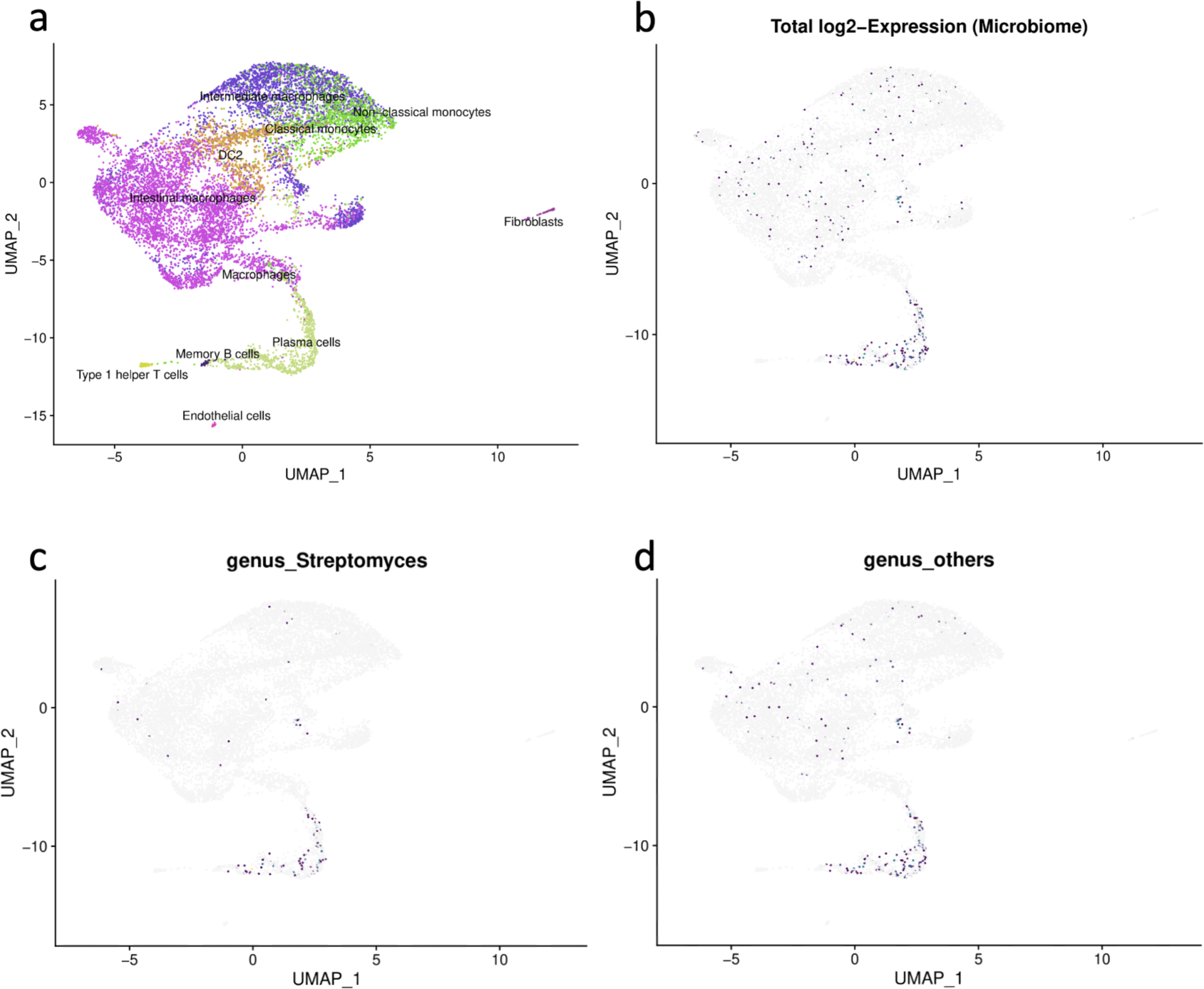
Cellsnake’s metagenomics feature was tested on mucosal macrophages. a) Four samples were integrated. The clusters were predicted and annotated using the CellTypist immune model. b) The cells annotated as “plasma cells” contain the highest number of bacterial reads. c) The foreign reads were mostly associated with Streptomyces. d) Cellsnake reports the top 10 most prevalent taxonomic groups by default. The rest collapsed and were reported as “others”. The user can select the desired taxonomic level (in this case, it was genus). All results are also saved as tables which include reads detected per cluster and annotation.

## Discussion

In recent years, there has been an increasing interest in scRNA-seq as it is a powerful technique for understanding the cellular heterogeneity of tissues and organs. However, the scRNA-seq data analysis can be complex and time-consuming. Cellsnake was designed to simplify this process, enabling researchers without extensive bioinformatics experience to easily analyze their data. It includes a range of automated preprocessing and downstream analysis tools and also provides advanced features for additional analysis. Its user-friendly interface and reproducibility features make it a valuable tool for researchers seeking to understand transcriptional heterogeneity in tissues at single-cell resolution.

Cellsnake has several critical functionalities for scRNA-seq data analysis. It includes preprocessing steps such as QC, filtering, and parameter auto-selection, and also has downstream analysis tools for identifying differentially expressed genes, performing clustering, visualization, and exploring cell type-specific gene expression patterns. These features are crucial for characterizing cell subpopulations and identifying specific genes and pathways associated with them. Cellsnake also includes advanced features such as supporting the integration of multiple scRNA-seq datasets to identify shared and unique cell types across different tissues or conditions. Cellsnake also ensures reproducibility by creating separate folders when required, restricting the versions of the tools in the environment, saving config files with the cellsnake version, explicitly sharing different images for each version in the Docker repository, and storing results for downstream analysis by default. In comparison to other tools (Table 3), cellsnake has several advantages, including a comprehensive range of tool utilization, unique features, the ability to run locally or on HPC platforms, and seamless integration with other workflows using Docker or Bioconda. Additionally, cellsnake also provides RDS files to enhance data sharing and accessibility.

**Table 3.** Standard features of cellsnake compared to available holistic tools/workflows.

|  | Cellsnake | Cellenics | Single Cell Omics Workbench |
| --- | --- | --- | --- |
| Platform | Snakemake/Python wrapper/Docker | Web based | Web based (Galaxy) |
| Input file type | Count tables (10X or others), R Data File | Count tables (10X) | Count tables (10X), FASTQ and others |
| Doublet filtering | Yes | Yes | No |
| MT gene filtering | Yes (auto) | Yes (auto) | Yes |
| Find clusters | Yes (auto) | Yes | Yes |
| Clustree plot | Yes | No | No |
| Differential expression analysis | Yes | Yes | Yes |
| Enrichment analysis | KEGG and GO | No | No |
| Celltype Annotation | Yes | Yes | No |
| Detailed gene expression plots | Yes | No | No |
| Metagenome analysis | Yes | No | No |
| Trajectory analysis | Yes | Yes | Yes |
| Integration | Yes (Seurat only) | Yes (various algorithms) | Yes |
| Output and downstream analysis | Plot files, expression tables, Seurat RDS files and excel files etc. | Plots and expression tables | Miscellaneous |

Recent studies have shown that the heterogeneity in microbiota and the present cell types along with their functions are co-dependent [28]. Cell-associated microbial reads can be identified in scRNA-seq data [29]. Cellsnake uses Kraken2 [24] to analyze this data and cellsnake provides the ability to fine-tune parameters to increase sensitivity and/or specificity and to use personal databases. This can help researchers identify potential microbial associations with host cells and tissues. Some of these microbial hits can originate from environmental contamination or can be false positives. These outcomes might not necessarily reflect real biological associations; nevertheless, the results may provide valuable insights for QC such as recognizing potential contamination sources.

There are some limitations of the workflow that need to be addressed. Firstly, cellsnake requires disk space to keep track of the entire pipeline, including metadata files that are required for advanced downstream analysis. Although the users can delete large files, they may want to keep metadata files for reproducing the results at a later time. Secondly, the fully-featured workflow relies on Cellranger outputs from 10x Genomics platform, which may not always be available. Even though cellsnake was designed and tested utilizing this platform, it can still use the count matrix files from other platforms, such as the fetal brain dataset. Third, while cellsnake has moderate performance in terms of memory and speed on standard workstations for an average number of cells, the auto-detection of parameters (e.g., resolution parameter) can be slow when processing samples with a large number of cells. To improve performance, a parallel version of the MultiK tool was used, which is not officially supported by the authors of MultiK (see materials and methods). Finally, the underlying tools utilized by cellsnake may involve various parameters. The fundamental parameters can be adjusted by the user and supplied through the configuration files, while the rest are set to default values. This approach was preferred to make the workflow more user-friendly.

In conclusion, cellsnake is a convenient and adaptable tool, empowering researchers to analyze scRNA-seq data in a reproducible and customizable manner. With its advanced features and streamlined workflow, cellsnake stands as a valuable bioinformatics asset for investigating cellular heterogeneity and gene expression patterns at single-cell resolution within tissues.

## Future Directions

Accurate bioinformatics software requires long-term development and commitment to the project [30]. It is also a major problem in the field that many projects are abandoned after publication, becoming unusable and outdated. For instance, cerebroApp [31], a component of cellsnake’s development version, was dropped as it is no longer in active development. Cellsnake is an open-source tool that is actively developed, allowing anyone to open pull requests and report issues on its GitHub page. To keep the software bug-free and streamlined, future developments of cellsnake will involve incorporating new tools, such as the latest Seurat version, and removing obsolete tools from the main workflow. The users can access the previous releases for reproducibility. Although cellsnake is mainly designed for the 10X Genomics single-cell platform, we plan to expand its compatibility with other platforms and offer additional support for various input formats. Our aim is for cellsnake to become an essential toolkit for fast, accurate, tunable, and comprehensive scRNA data analysis.

## Data availability

The publicly available datasets for the fetal brain and liver are available under accessions PRJNA429950 and PRJEB34784, respectively. Macrophage-only samples from gut mucosal tissue are deposited in the European Genome-Phenome Archive (EGA) under the following accession numbers: EGAD00001007765 and EGAS00001005377. The EGA deposited files are under controlled access, requiring the data access committee permission for retrieval. The cellsnake analysis results on test samples are available at https://doi.org/10.5281/zenodo.8282676. A copy of the fetal brain dataset can also be found in our frozen Zenodo repository.

## Availability and Requirements

- Project name: cellsnake
- Project homepage: https://github.com/sinanugur/cellsnake
- Documentation: https://cellsnake.readthedocs.io/en/latest/
- RRID: SCR_023666
- Operating system: Platform independent
- Programming language: Python, R
- Other requirements: Python 3.8 or higher, R 4.2.2
- License: MIT
- PyPi: https://pypi.org/project/cellsnake
- Bioconda: https://anaconda.org/bioconda/cellsnake
- Docker: https://hub.docker.com/r/sinanugur/cellsnake
- Snakemake workflow: https://github.com/sinanugur/scrna-workflow

## Author contributions

SUU devised the project, created the workflow, wrapper, and R scripts, and drafted the manuscript with input from all authors. KRV contributed to the figures and the R scripts. VTK contributed to the R scripts. MC contributed to the R scripts and revised the preliminary manuscript. ESB revised the manuscript and acquired financial support. FLJ revised the manuscript and acquired financial support. DD supervised the project, contributed to the R scripts, revised the manuscript, and acquired financial support.

