## supplementary figures for "cellsnake: a user-friendly tool for single-cell RNA sequencing analysis"

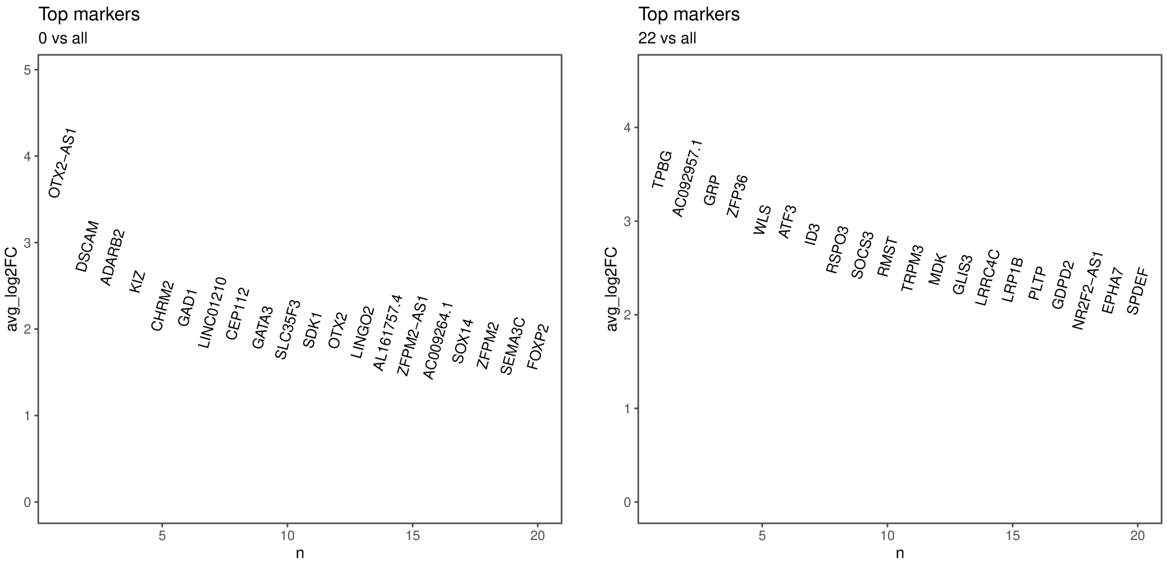


**Supplementary figure 1.** Summarized marker plots show the top 20 genes for each cluster predicted which gives an overview of the cluster markers. Here we show upregulated genes in the first and the last cluster predicted. These results are based on the integrated fetal brain dataset.

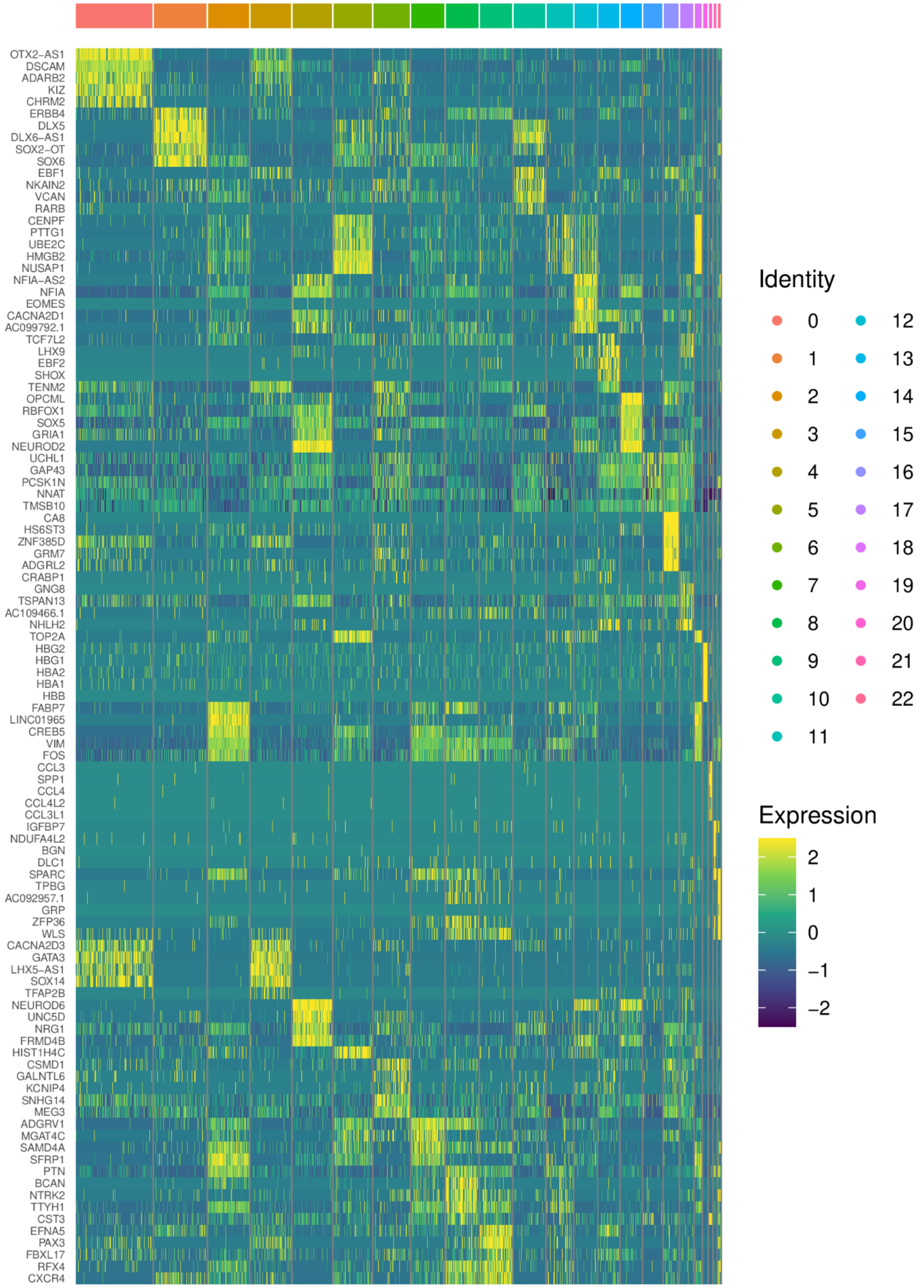


**Supplementary figure 2.** The heatmap plots provide another way of cluster marker visualization. The clusters are based on the integrated fetal brain dataset.

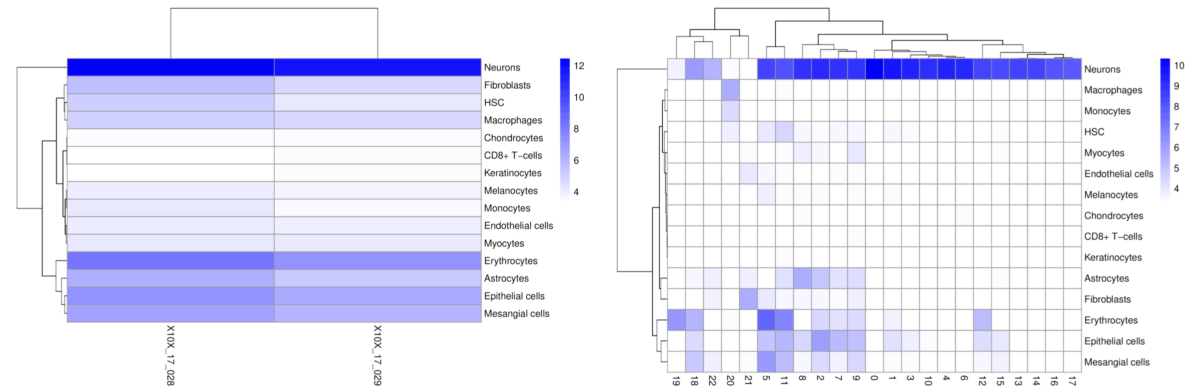


**Supplementary figure 3.** In addition to generating dimension reduction plots with annotations, the SingleR package allows for obtaining annotation scores based on clusters. The left panel demonstrates the ability to compare annotation scores of samples within an integrated object. Specifically, in the displayed example, two samples are derived from the fetal brain dataset.

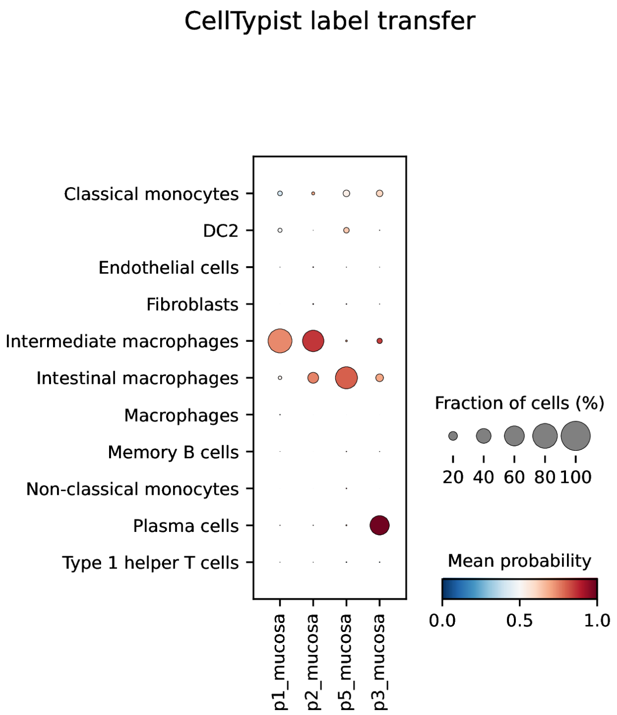


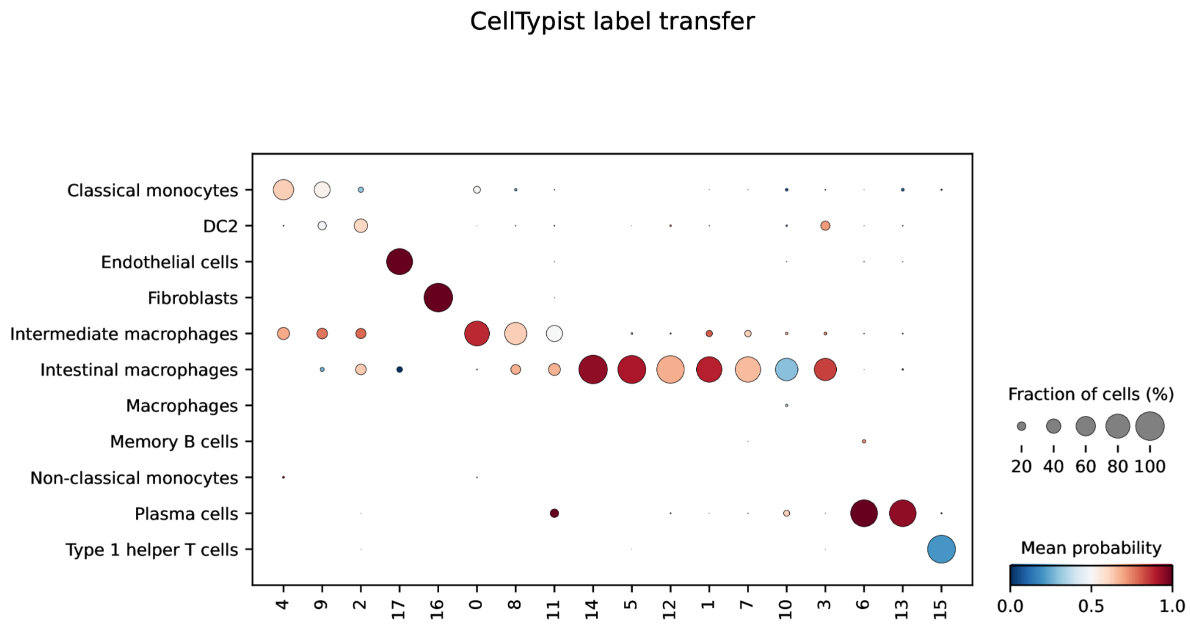


**Supplementary figure 4.** In addition to the SingleR package, cellsnake incorporates CellTypist for label transfer and annotation purposes. CellTypist enables the generation of probability scores at both the cluster and sample level. To illustrate this functionality, we provide an example using the mucosal macrophages dataset.

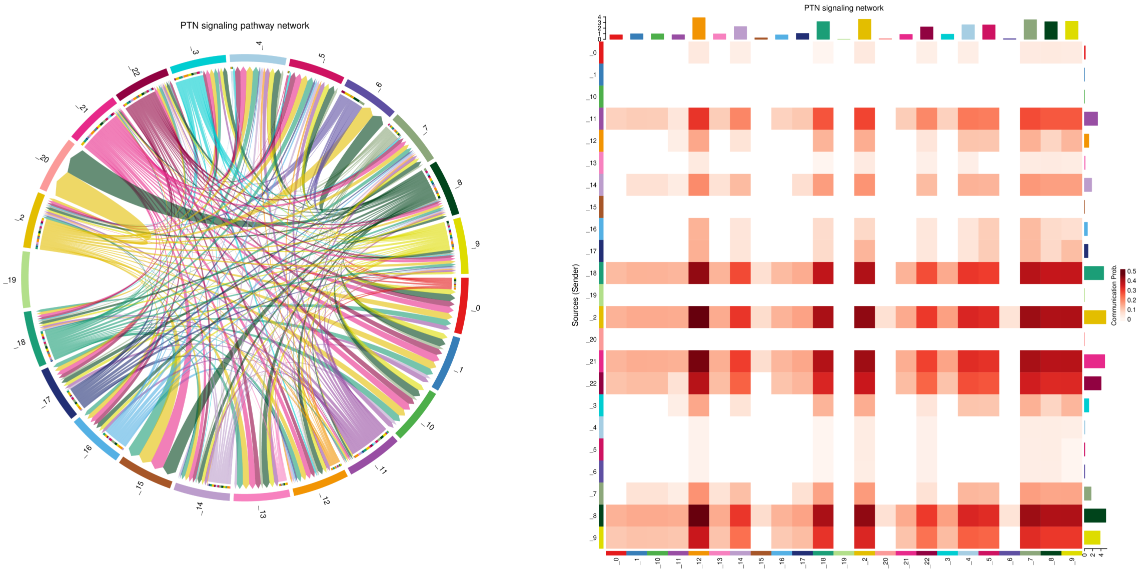


**Supplementary figure 5.** CellChat provides significant signaling pathway visualization among predicted clusters. For instance, the PTN signaling pathway is exemplified here, with the clusters being derived from the integrated fetal brain dataset.

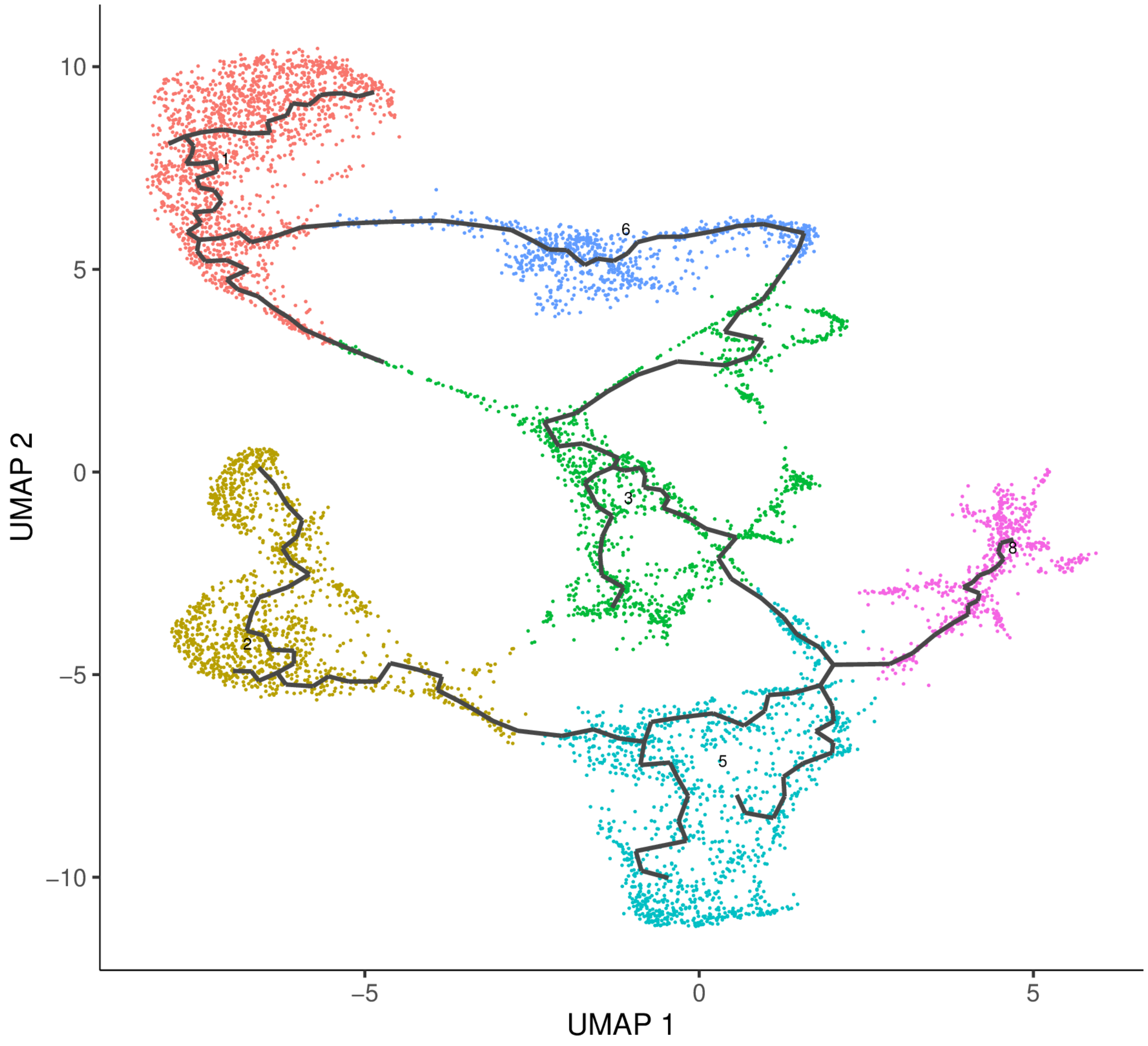


**Supplementary figure 6.** Cellsnake uses Monocle 3 for trajectory analysis. Here the result is based on the integrated fetal brain dataset, partition 1.
